# Electron beam lithography fabrication of SU-8 polymer structures for cell studies

**DOI:** 10.1101/849745

**Authors:** Jakob Vinje, Kai S. Beckwith, Pawel Sikorski

## Abstract

Flat surfaces decorated with micro- and nanostructures are important tools in biomedical research used to control cellular shape, in studies of mechanotransduction, membrane mechanics, cell migration and cellular interactions with nanostructured surfaces. Existing methods to fabricate surface-bound nanostructures are typically limited either by resolution, aspect ratio or throughput. In this work, we explore electron beam lithography based structuring of the epoxy resist SU-8 on glass substrate. We focus on a systematic investigation of the process parameters and determine limits of the fabrication process, both in terms of spatial resolution, structure aspect ration and fabrication throughput. The described approach is capable of producing high-aspect ratio, surface bound nanostructures with height ranging from 100 nm to 4000 nm and with in-plane resolution below 100 nm directly on a transparent substrate. Fabricated nanostructured surfaces can be integrated with common techniques for biomedical research, such as high numerical aperture optical microscopy. Further more, we show how the described approach can be used to make nanostructures with multiple heights on the same surface, something which is not readily achievable using alternative fabrication approaches. Our research paves an alternative way of manufacturing nanostructured surfaces with applications in life science research.

## 1 Introduction

Nanotechnology has in the recent years greatly increased our ability to control and study matter on the length-scales between 1 and few hundreds nanometers. This increased control of materials has been important for fabrication of electronic circuits with smaller component size,^1^ in production of advanced catalysts nanomaterials^2, 3^ and for nano-particle use in photovoltaics.^4, 5^ These development in nanofabrication have been supported by advances in characterisation techniques allowing for wider areas of application.

In the field of biomedical research, nanotechnology has been seen as a promising technology to develop new research tools, new diagnostic devices or even novel drug delivery strategies. This is due to the ability to structure surfaces or fabricate free standing structures like nanoparticles with a resolution comparable to the size of sub-cellular compartments and organelles. For example, Cai *et al.* used nanofabrication based on electron beam lithography (EBL) to define gold structures on a glass surfaces to precisely control position receptor ligands and to subject T cells to defined arrangements of ligands.^6^ This is due to the fact that T cell receptor clustering by surface-bound ligands is believed to be a highly effective method to triggering the tyrosine kinase cascade leading to T cell activation.^7^

Another type of nanostructured cell culture substrates that has recently attracted a lot of attention, are based on flat and cell compatible supports decorated with vertically aligned nanowires or nanopillars.^8–12^ Nanowires or nanopillars are typically 10 nm to 100 nm in diameter 1 µm to 5 µm in length and can be made from carbon,^13, 14^ SiO, silicon,^15, 16^ indium arsenide^17^ or copper oxide.^18^ Fabrication is typically performed using a combination of patterning (UV or EBL), growth (for example molecular beam epitaxy, plasma-enhanced chemical vapour deposition and electrodeposition) or etching (by wet- or dry-etching techniques). Such substrates have been used to deliver biologically relevant molecules into cells, monitoring enzyme activity in living cells, and to study cell viability, adhesion and morphology on in response to nanowire geometry.

Amin et al. used arrays of vertical nanopillars functionalised with a adhesion-promoting molecule for selective guidance of primary hippocampal neurons.^19^ A fabrication strategy based on an electron sensitive polymer resist deposited onto a silicon nitrate membrane was used. With energy from secondary electrons from ion beam milling, nanopillars with height of 1.8 µm and diameter of 150 nm could be fabricated

Gopal et al. used porous silicon nanoneedles to study the delivery of biological payloads in human mesenchymal stem cells.^20^ For substrate fabrication, the authors use a combination of UV lithography, metal-assisted chemical etching and reactive ion etching. Cellular processes in human mesenchymal stem cells (hMSCs) were studied using a combination of optical and FIB-SEM microscopy, scanning ion conductance microscopy and molecular biology techniques. They were able to show that that payload delivery to cells in culture is at least partly mediated by endocytic processes which were observed to be upregulated selectively at the cell-nanoneedle interface.

Realisation of many of nanostructured cell culture substrates require fabrication methods that allow to produce structures with an arbitrary shape and with resolution of at least 100 nm. Devices fabricated for biomedical applications require the active area decorated with nanostructures to be in the range of *>* mm^2^. This is due to the physical size of cells and a need to measure sufficient number of events to conclude about processes in the studied system. It is important that both the nanostructures and the surrounding substrate are bio-compatible and stable under physiological conditions. It is also necessary that the nanostructured substrates can be integrated with other techniques commonly used in the life sciences research, including optical microscopy, well-based assays, microfluidic systems or biosensors.^21^ In addition, many biomedical applications require single use, disposable substrates. As a result employed nanofabrication methods should have sufficiently high throughput.

Micro- and nanostructured surfaces can be fabricated using various lithographic techniques. Fabrication approaches such as mask-based photolithography^22^ and mask-less photolithography^23, 24^ are frequently applied, but structure resolution are for both techniques limited by the optical diffraction limit. Other approaches such as dip pen lithography,^25^ nano-sphere lithography,^26^ block-copolymer lithography^27^ and other similar approaches^28^ are able to perform structuring with sub-100 nm resolution, but have a limited throughput.

Electron beam lithography (EBL) is able to bridge the gap between high throughput and the high spatial resolution patterning techniques. EBL relies on scanning a electron beam across a surface covered with a film of a electron sensitive resist. Energy transfer from the electron beam induces chemical reactions in the resist, which locally changes its solubility. Depending on the type of resist, either exposed (negative tone resist) or unexposed areas (positive tone resist) can be dissolved during the development process. For both resist types, a minimal critical dose (energy per area) is needed for the chemical reactions to be induced to a sufficient degree. For some negative resists, irradiation with electrons generates a chemical catalyst initiating a number of cross-linking/polymerisation reactions when heated. In a EBL fabrication process, a computer designed mask defines exposed areas, and arbitrary structures within the resolution limit may be fabricated.

In EBL, it is the scattering of electrons in the resist that determines the ultimate resolution of the fabrication process. Even though the diameter of the primary electron beam can be as small as below 1 nm, the scattering events will cause the electron beam to broaden, as the electrons travel through the resist film. This broadening makes the resolution dependent on the resist film thickness and limits minimum size of fabricated structures. Additionally, some of the electrons from the electron beam might scatter at large angles in the resist or substrate and can travel for approximately 30 µm in the horizontal direction.^29^ As a consequence, the electrons are not entirely confined to the exposed areas but might also deposit energy in adjacent areas, known as proximity effect.

For high resolution EBL these two effects influence the fabrication outcome. If exposed with the same dose, isolated features of the pattern may be underexposed, whereas denser areas of the pattern might be overexposed. To ensure correct pattern replication an proximity effect correction (PEC) has to be performed. By numerically estimating the contribution from the proximity effect, the dose throughout the pattern can be adjusted, such that the energy deposition from adjacent areas of the pattern can be accounted for.

Fabrication of surface bound nanostructures with EBL can be done by spin coating of a wide range of electron sensitive polymers on different flat substrates such as silicon or glass. One candidate polymer is the epoxy based resist SU-8.^30, 31^ SU-8 was originally developed for UV-photolithography, but is also sensitive to electron exposure.^32^ It can be applied to substrates in a wide variety of thicknesses, ranging from more than 300 µm down to sub-micrometer thickness (MicroChem datasheet/processing guidelines). This versatility has made it a frequently used resist in production of microelectromechanical systems (MEMS), as moulds for making stamps in soft lithography and as mould for fabrication of microfluidic devices.^33–35^

SU-8, is a chemically amplified resist, with resin monomers (glycidyl ether derivative of bisphenol-A novolac), an organic solvent (cyclopentanone) and a photo-acid generator (triarylsulfonium hexafluoroantimonate salts). Upon exposure, the photo-acid generator is activated creating a strong acid. This acid catalyses the reaction forming a highly cross-linked network upon subsequent baking.^36^ Diffusion of the photo-acid generator contributes to a general broadening of the exposed structures.^37, 38^ SU-8 is stiff with a Young’s modulus reported in the range from 0.9 GPa to 7.4 GPa,^32, 39^ its processing is well established,^40^ it has high chemical stability^39^ and it is known to be bio-compatible.^41^ High resolution, high aspect ratio combined with rapid fabrication time due to very high sensitivity^42^ makes SU-8 EBL fabrication a promising approach for further exploration as structural component in structured cellular substrates.^11, 43^ Furthermore, SU-8 is suitable for grey-scale electron beam lithography due to its low contrast.^44^ Fabrication of high aspect ratio structures using a 100 keV beam has previously been performed, realising structures with heights up to 5 µm.^30^ Bilenberg *et al.* tested the resolution for nanofabrication of SU-8 using a 100 keV EBL system and showed that lines with widths of 24 nm can be written in a 99 nm thick resist.^44^

In previous work,^11, 43^ our focus has been on exploring cell behaviour on substrates decorated with SU-8 structures, however in this work we focus on exploring the limitations to our fabrication process to understand which structures we are able to fabricate in this system. In this work we present a flexible approach for high-throughput fabrication of SU-8 nanostructures on glass using electron beam lithography. To test the limitations of the fabrication process, we focus on two test-patterns; arrays of lines with variable line width and variable spacing between each line, and nano-pillar arrays with variable pillar-spacing. We analyse fabrication outcomes for resist thickness in the range between 25 nm to 8000 nm. For each thickness and test pattern type (lines and pillars), the electron exposure dose that produces the best fabrication outcome is presented. Finally, we show how adaptation of the fabrication process can be used to make structures of two different heights on the same substrate.

## 2 Experimental Section

### 2.1 Materials and Methods

#### 2.1.1 SU-8 resist

SU-8 2010 (MicroChem Corp.) was used as a base resist to prepare films in a range between 25 nm to 8000 nm. The base resist, SU-8 2010 is designed to give film thickness of 10 000 nm when spin coated at 3000 rpm. According to manufacturer specifications it can be diluted using cyclopentanone resulting in reduced film thickness at the same spin speed. Based on the data-sheet provided by the manufacturer, dilutions listed in Table 1 were prepared. Spin curves for all dilutions are shown in Figure 1. These were acquired by spin coating different SU-8 dilutions on silicon wafers at velocities ranging from 1000 to 6000rpm. Film thickness was measured using a F20 Reflectometer (Filmetrics) at a minimum of three different positions on the sample. Data from the spin curves was used to select process parameters for making SU-8 films with desired thickness. The solutions were stored in resist bottles (amber glass bottles with polypropylen lid and low-density polyethylene insert, Rixius AG) at room temperature.

**Figure 1:**
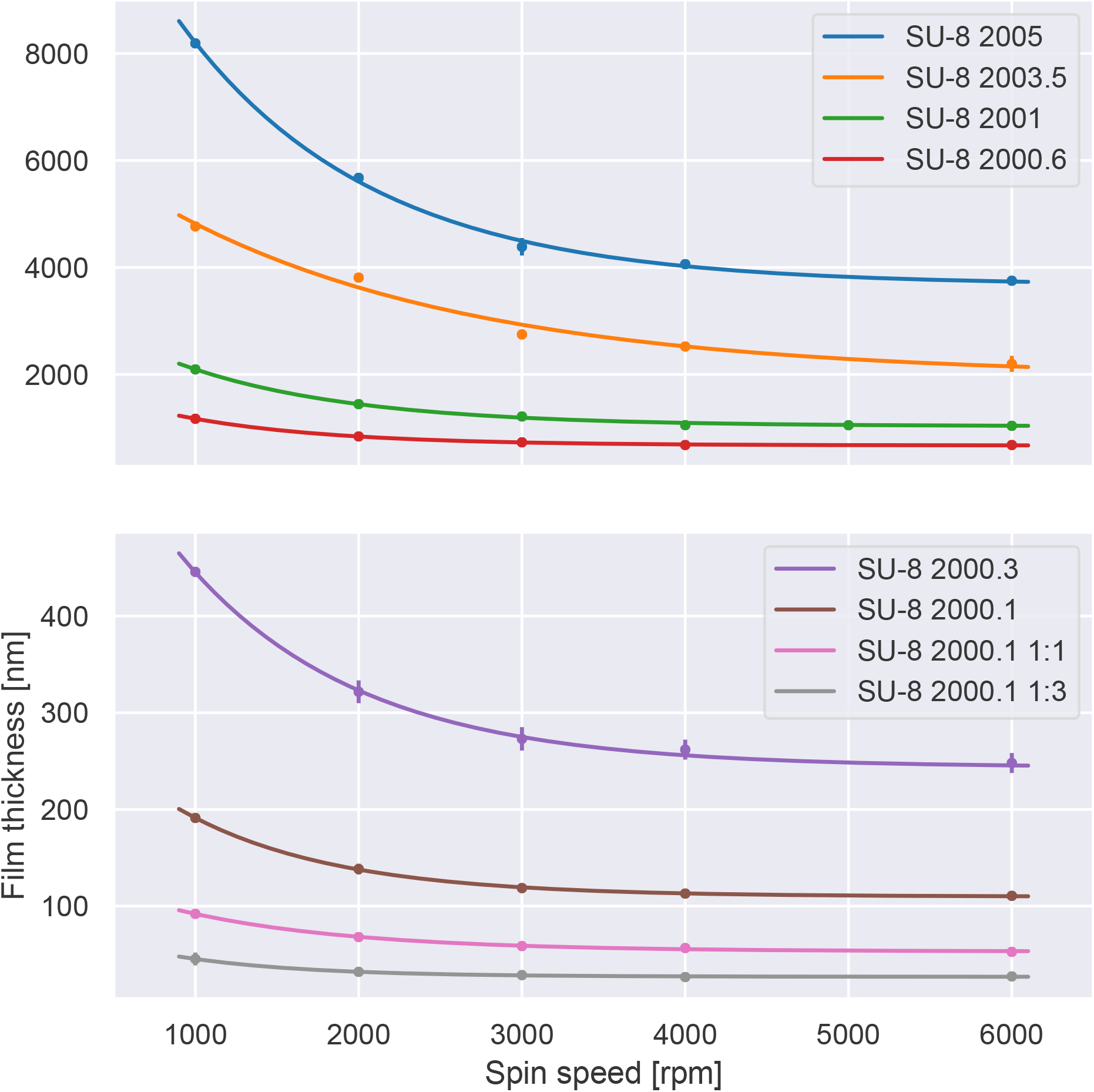
Thickness of spin coated SU-8 dilutions measured on HMDS treated silicon substrates. All thicknesses were measured minimum three times on two separate samples. To each of the dilutions, an exponential curve is fitted. With these dilutions, it is possible to spin coat SU-8 with thicknesses ranging from approximately 8000 nm down to 25 nm. By varying the solid content, the viscosity properties of SU-8 can be controlled to produce films with a range of thicknesses in this range.

**Table 1:** Formulations of SU-8 diluted from SU-8 2010. SU-8 2010 has a solid resin content of (58%), and formulations with lower solid content was prepared by adding the resist thinner cyclopentanone. Formulation names was created based on the naming from the manufacturer, corresponding to the approximate thickness of a film when spin coated at 3000rpm.

| Formulation name | Approx. Thick. 3000 rpm [nm] | Solid content |
| --- | --- | --- |
| SU-8 2005 | 5000 | 45% |
| SU-8 2003.5 | 3500 | 37% |
| SU-8 2001 | 1000 | 25% |
| SU-8 2000.6 | 600 | 18% |
| SU-8 2000.3 | 300 | 9% |
| SU-8 2000.1 | 100 | 4% |
| SU-8 2000.1 1:1 | 50 | 2% |
| SU-8 2000.1 1:3 | 25 | 1% |

#### 2.1.2 Sample preparation for EBL

24 mm × 24 mm glass cover slips (#1.5, Menzel-Gläser, 170 µm) were cleaned by immersion in acetone, isopropanol, and rinsed in de-ionised water followed by 2 min oxygen plasma treatment (Diener Femto plasma cleaner, power 100 W and pressure 0.4 mbar). The substrates were then dehydrated at 150 *^◦^*C on a hotplate for 10 minutes. For some samples, hexamethyldisilazane (HMDS) was used as a adhesion promoter (MicroChem Corp. SU-8 2000 Processing Guidelines). It was applied by placing dehydrated glass substrates in a desiccator connected to a diaphragm pump and containing a vial of HMDS (Product Number: 440191, Sigma Aldrich). The sample was then kept under HMDS atmosphere for 60 min. Spin coating of the resist was performed straight after HMDS treatment.

Based on the spin curves, (Table 1 and Figure 1) films with thickness of 25 nm, 50 nm, 100 nm, 250 nm, 500 nm, 1000 nm, 2000 nm, 4000 nm and 8000 nm were made on glass cover slips. All films were then inspected with F20 Reflectometer (Filmetrics) to verify thickness and by an optical microscope to verify the film quality. Three measurements were taken at different locations to check for film uniformity. Soft bake was performed on a hotplate at 95 *^◦^*C for a duration given in Figure 2. Soft bake time was found by interpolating the data from the processing guidelines (URL: http://microchem.com/pdf/SU-82000DataSheet2000_5thru2015Ver4.pdf). A 50 nm layer of conductive polymer AR-PC 5091 Electra 92 (AllResist GmbH) was spin coated (2000 rpm for 60 s) on top of the SU-8.

**Figure 2:**
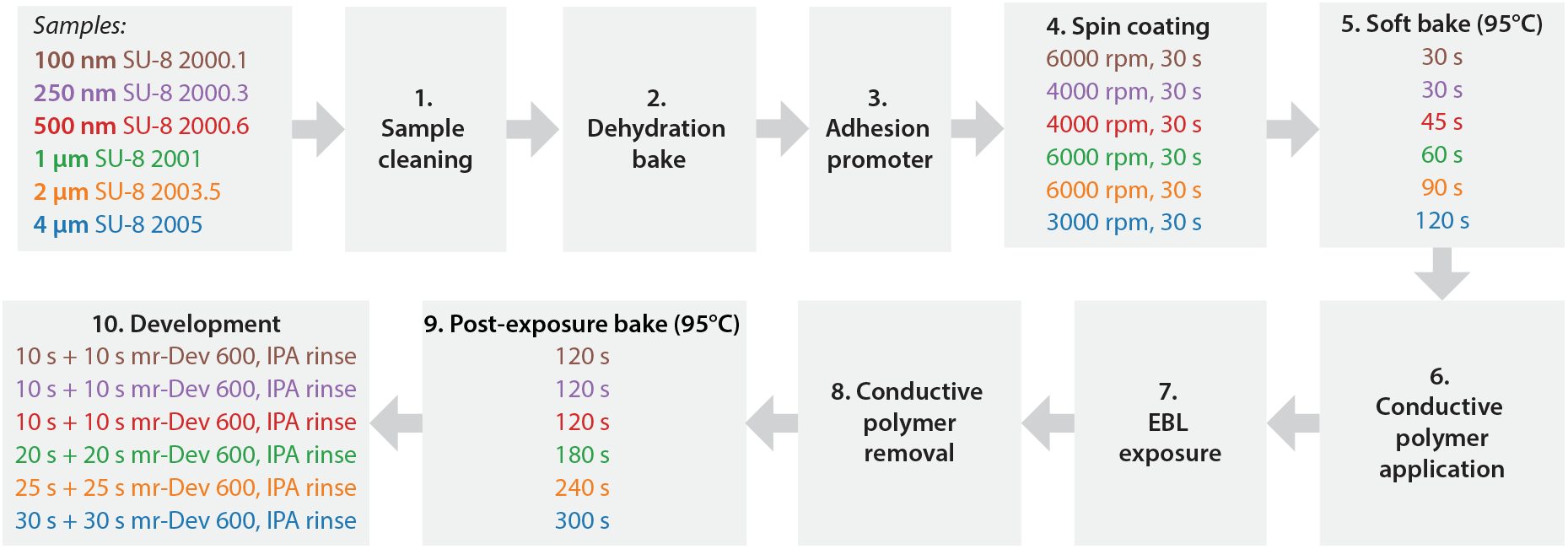
Processing of SU-8 films on glass substrates. The processing starts leftmost in the figure (step 1) and progresses through each of the steps up to 10. Steps 1,2,3,6 and 8 are equal for all samples and is described in detail in the text. Step 7, the EBL exposure, is also described in this section.

#### 2.1.3 Pattern design

Two types of exposure pattern designs were created: line patterns with variable line width and line separation and dot patterns that result in a nanopillar topography.

Line patters were designed using a LayoutEditor (build: 20190401, Juspertor GmbH). Before exposure, proximity effect correction was performed using using a commercially available proximity correction software (Beamer version 5.6.2, Genisys GmbH) and exported to a file containing information about design geometries and the proximity effect corrected dose at each location in the design readable by the Elionix EBL system. Table 2 lists all designed combinations of line widths and separations. Each pattern was written on a area of 250 µm by 250 µm. For nanopillar arrays, the built-in dot-pattern generator in Elionix WeCAS CAD-software was used to create hexagonal pillar patterns with pillar to pillar distances (array pitch) of 100 nm, 250 nm, 500 nm, 750 nm, 1000 nm, 2000 nm, 5000 nm and 10 000 nm. Each pillar array was written covering an area of 500 µm by 500 µm. Line pattern, pillar pattern and relevant array parameters describing the geometry are schematically shown in Figure 3.

**Figure 3:**
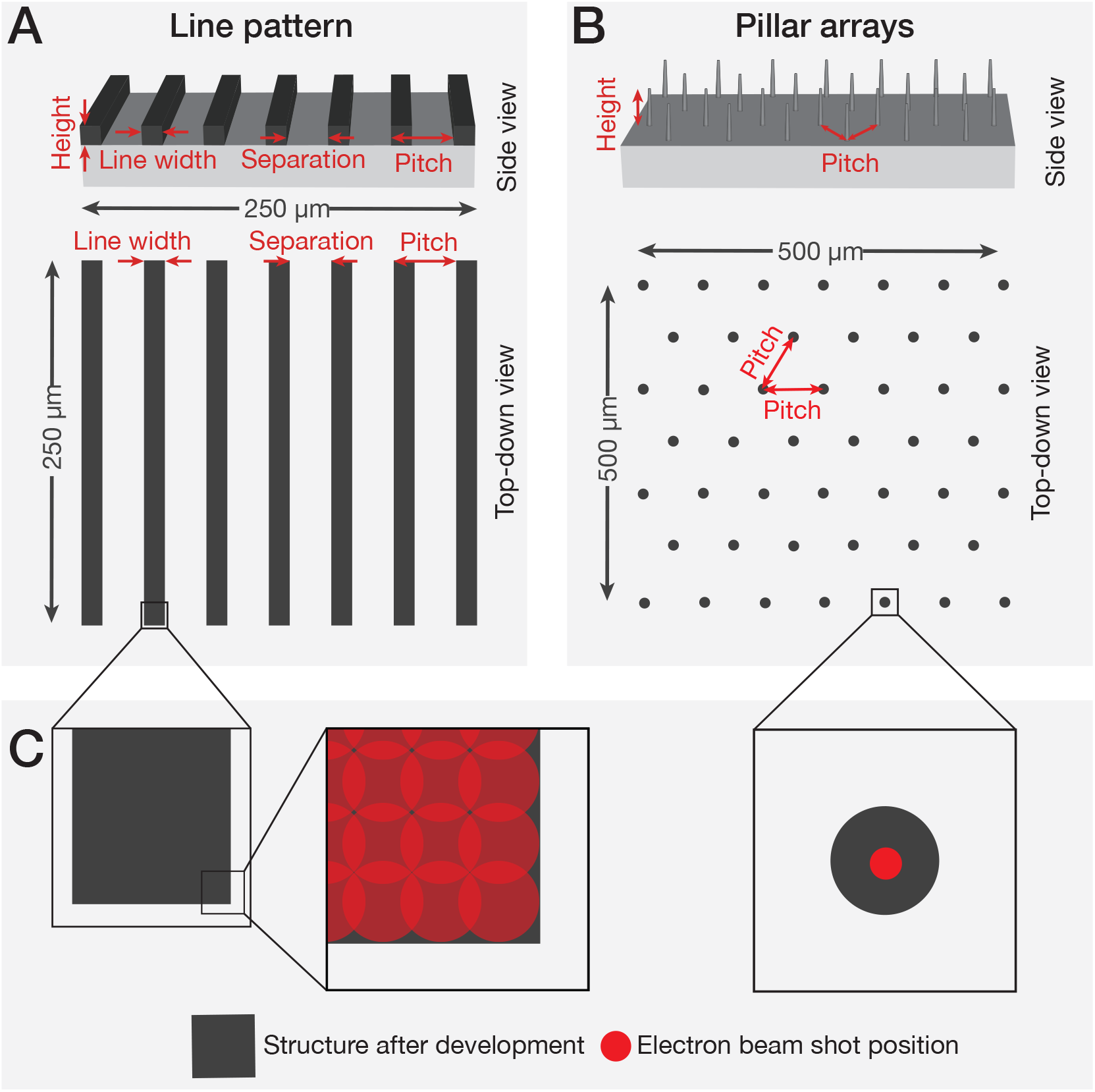
A) Schematic illustration of line-pattern and relevant sizes from a top down and side-view. The pattern is replicated over an area of 250 µm by 250 µm. B) Top down and side-view of fabricated pillar structures. The exposures are set up by defining a grid of dots and corresponding dose at each of these dots. C) Detailed view of part of line pattern and single pillar. Red circles indicate exposure positions. Exposure is done with overlapping exposure positions, with a low exposure time per position. This gives a continuous structure after development. Pillars are exposed as single dots by keeping the beam stationary and exposing one shot for an extended amount of time. The figures are for reasons of illustration not drawn to scale.

**Figure 4:**
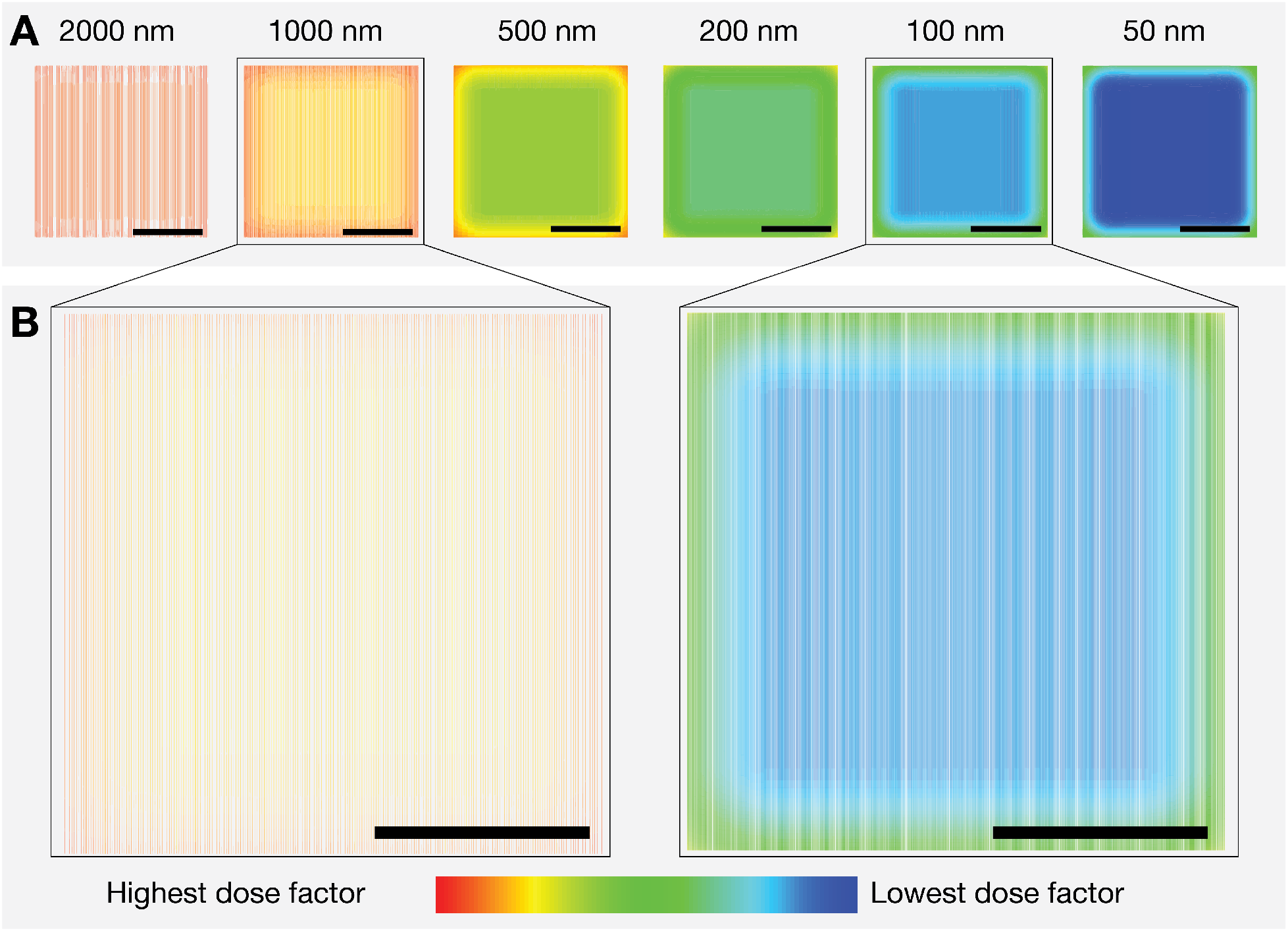
Outcome of BEAMER simulation showing the calculated dose coefficient for line pattern areas of 100 nm wide lines in the line arrays. The calculated dose coefficient is used to adjust the “base dose” set when performing an exposure. A) Calculated dose coefficient for arrays of 100 nm parallel lines with variable separation between each line. The lines with highest separation are calculated to receive a higher dose than the denser patterns with lower line to line separation. B) Detailed view on line patterns with 1000 nm and 100 nm line spacing. For both patterns calculations indicate that the centre of each line pattern receives a comparably lower dose compared to the edges. This effect can be attributed to the proximity effect.

**Table 2:** Tested Line-width and Separation Combinations. The tests were done on all film thicknesses.

| Line width [nm] | Separation between lines [nm] |  |  |  |  |  |
| --- | --- | --- | --- | --- | --- | --- |
|  | 20 | 10 | 5 | 2 | 1 | 0.5 |
| 1000 | — | 10000 | 5000 | 2000 | 1000 | 500 |
| 500 | — | 5000 | 2500 | 1000 | 500 | 250 |
| 250 | — | 2500 | 1250 | 500 | 250 | 125 |
| 100 | 2000 | 1000 | 500 | 200 | 100 | 50 |
| 50 | 1000 | 500 | 250 | 100 | 50 | 25 |
| 25 | 500 | 250 | 125 | 50 | 25 | 12.5 |

#### 2.1.4 Electron beam lithography

Elionix ELS-G100 EBL-system, operating at 100 kV acceleration voltage was used to structure SU8 films. Glass cover slips were mounted on a multi-piece sample holder, with each sample placed in a L-shaped sample slot and secured with a clamp. For each exposure, two samples could be loaded and exposed.

Line patterns were defined using direct writing, vector scanning mode, with a current of 200 pA (resit thicknesses of 100 nm, 250 nm and 500 nm) or 500 pA (resist thicknesses 1000 nm, 2000 nm and 4000 nm) and an beam size of *≈* 2 nm. All exposures were done with 500 µm write field size. Structures in one write field are written without moving the sample stage, just by deflecting the electron beam.

Exposing areas in the EBL requires a base dose to be chosen. The base dose is then adjusted with a dose factor found when performing proximity effect correction (PEC). A line pattern dose test was performed by repeating the pattern six times with a 50% increase in base dose for each repetition. For thin SU-8 films (resit thicknesses of 100 nm, 250 nm and 500 nm), a starting base dose of 2 µC cm*^−^*^2^ was selected, for thicker films (resist thicknesses 1000 nm, 2000 nm and 4000 nm) a starting base dose of 7 µC cm*^−^*^2^ was selected. PEC was performed for each film thickness by applying a dose correction factor to the set base dose to all locations in the pattern. The dose was adjusted with a factor between 0.86 and 1.69.

All pillar arrays were exposed using built in dot-pattern generator (DPG) in the Elionix EBL software with a beam current of 500 pA and a beam size of *≈* 2 nm. Arrays were exposed over an area of 500 µm by 500 µm, all within one write-field with the same size. The exposure dose for pillar patterns was then optimised by performing a dose test. The dose range for dot-exposures were set up such that each array with a given pitch had a starting dose per dot and the dose was then increased by 25% per repetition of that array. Each array was then repeated 8 times for each film thickness. Based on preliminary experiments a starting dose per dot for dot arrays were set as specified in Figure 9.

After exposure, samples were rinsed in de-ionised water for 60 s to remove AR-PC 5091 Electra 92 conductive polymer and then dried with nitrogen stream. Samples were post-exposure baked at 95 *^◦^*C on a hotplate for a duration specified in Figure 2. Post-exposure bake times were based on the processing guidelines supplied by the resist manufacturer (URL: http://microchem.com/pdf/SU-82000DataSheet2000_5thru2015Ver4.pdf). The samples were developed twice in mr-Dev 600 developer (Micro resist technology GmbH) for times specified in Figure 2, rinsed in isopropanol and finally dried using nitrogen.

#### 2.1.5 Topographies with multiple heights

Samples with multiple structure heights could be fabricated with the same approaches as used for single-layer exposures. The first layer of SU-8 structures were fabricated as previously described above (steps 1 to 9 in Figure 2), but without performing any development. An additional layer could then be spin coated on top of the exposed, but not developed first SU-8 film. The conductive polymer was then spin coated on top of the SU-8 and structures were exposed in both structures. PEB and development was then performed with parameters corresponding to the total layer thickness.

#### 2.1.6 Sample morphology

Before scanning electron microscopy (SEM) samples were sputter coated with 5 nm Platinum/Palladium alloy using a 208 HR B sputter coater (Cressington Scientific Instruments UK) and mounted on standard SEM specimen stubs. Scanning electron microscopy was performed using a FEI Apreo SEM and an in-lens detector or an ETD detector. Imaging at normal incidence angle was performed using 2 kV acceleration voltage and 0.2 nA current, whereas the tilted imaging was done using 5 kV acceleration voltage and 0.2 nA current with the sample mounted on a 45° pre-titled stage with additional tilting of 30°. Normal incidence imaging was performed at a working distance of 3 mm, tilted imaging at a working distance of 10 mm.

### 2.2 Results

From the range of applications of nanostructured surfaces for cell studies presented previously, it is clear that different geometries and sizes are of use when interfacing cells and nanostructures. To a large extent these structures can be made with the epoxy based polymer using SU-8 resist and EBL and thus, our aim is to explore limitations of EBL fabrication process in SU-8 on glass substrates. To achieve this, we investigate how resist thickness and electron dose effect structure geometry. We provide design criteria and process parameters for making arbitrary structures withing the accessible shape range. Using dilutions of SU-8 listed in Table 1 we were able to produce uniform SU-8 films with thicknesses between 25 nm and 8000 nm. For all resist thicknesses, we investigate optimal dose, minimum size and minimum separation for fabrication of nanopillars, as well as optimal dose, minimum size and minimum separation for 2D structures modelled by line patterns. The line pattern consist of parallel lines, with different line widths and separations between lines, as described in Table 2. For nanopillar samples, hexagonal arrays with variable pillar to pillar spacing (pitch) in a range between 250 nm to 10 000 nm were made.

Successful fabrication requires optimisation of the writing dose, as a low dose results in incomplete cross-linking of SU-8 and structure instability or detachment. Too high dose results in loss of resolution and possibly cross-linking of the resist in the areas between fabricated structures. Fabrication of high resolution test structures was not possible in resists thicknesses below 100 nm or above 4000 nm. For the 50 nm resist, no clear exposed pattern was observed, only partly cross-linked SU-8 residues covering the exposed area and the close proximity. For resist with a thickness above 4000 nm, our test pattern was either exposed with a too low dose, completely cross-linked or had peeled off the substrate.

To mitigate charging problems, a 50 nm layer of conductive polymer was added on top of the SU-8 structures. After the EBL exposure, the polymer could be rinsed away in deionised water, without leaving any visible residue. Sample inspection after development revealed that the conductive layer prevented charging problems without causing any noticeable changes to exposure parameters such as the necessary electron dose.

Dose testing for the line patterns was performed by writing multiple copies of the same pattern on the same substrate. For thin SU-8 films (100 nm, 250 nm and 500 nm) a dose between 2 µC cm*^−^*^2^ and 22.8 µC cm*^−^*^2^ was tested. For thicker films (1000 nm, 2000 nm and 4000 nm) the dose range was between 7 µC cm*^−^*^2^ and 79.3 µC cm*^−^*^2^. In addition to the dose test, we performed dose proximity effect correction (PEC) that accounts for contributions from back scattering electrons from the nearby structures to the total dose (see M&M). After development, the surface morphology was investigated with SEM. Fabrication outcome was categorised based on the surface morphology as (1) free standing lines, (2) partly collapsed lines, (3) partial cross-linking of the resist in between the line, (4) complete cross-linking of the resist film. Figure 5 shows the outcome written with the optimal dose, for each combination of line width and line separation. The optimal dose for the pattern is selected as the lowest dose that gives sufficiently cross-linked SU-8 in the exposed regions. Using a dose above the optimal dose resulted in the loss of resolution both in terms of feature size and minimum separation between structures.

**Figure 5:**
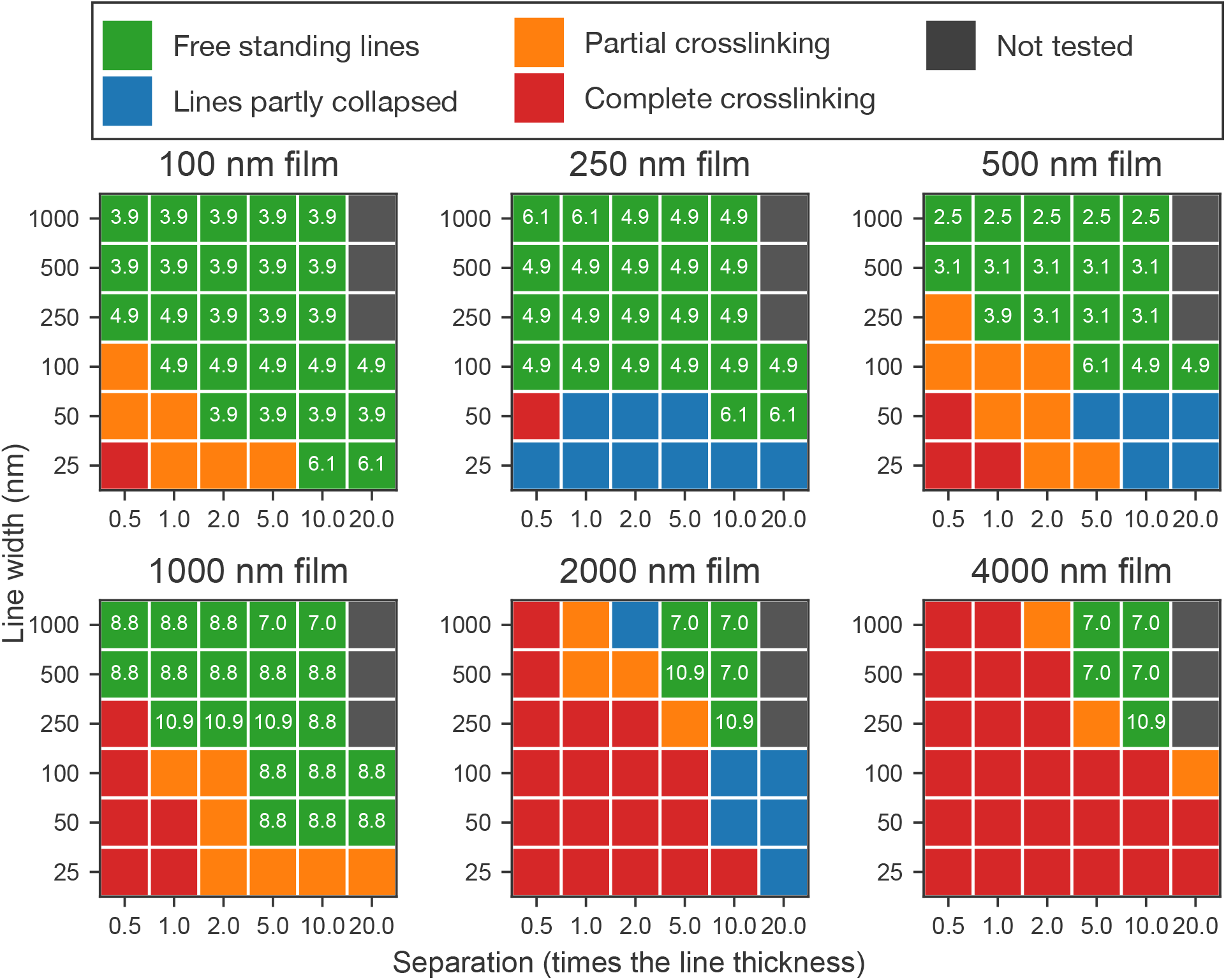
Colour coded representation of line pattern fabrication outcomes in SU-8 with different line widths and variable separations for various film thicknesses. The separation is given in multiples of the line width. Outcome is categorised following the different fabrication outcomes indicated in Figure 6. When spaced too closely together, exposed lines completely cross-link (indicated in red) and a continuous film over the entire array is formed. When the separation is increased, the contrast in received dose between exposed lines and gaps increases and free standing lines are formed (green colour). In-between these two regimes, partly cross-linked structures are observed. For given combinations of line width and structure height, only fragile, collapsed lines could be fabricated. For lines with width 250 nm, 500 nm and 1000 nm, 20 times separation was not tested. In the cases of successful fabrication of free standing lines, the electron dose producing the outcome is noted.

One challenge in successful fabrication of high aspect ratio lines is to ensure that fabricated lines remain intact and attached to the substrate after development. As described by Tanaka et al. pattern collapse can be classified into “deformation” or “peeling”, related to the mechanical stability of the remaining resist material and to the adhesion between the resist and substrate respectively.^45^ This is typically reported to occur when the sample is dried after the development step. For certain combinations of line width and separations, we also observe resist collapse, as shown in Figure 6C and G. This effect is primarily seen for closely spaced high aspect ratio lines, such as 25 nm thin lines in 250 nm SU-8 films. Collapse due to lack of mechanical stability in the resist is the predominant mechanism for line structures. As seen from Figure 7 narrow lines remained fragile even at higher separations, whereas the wider lines were stable.

**Figure 6:**
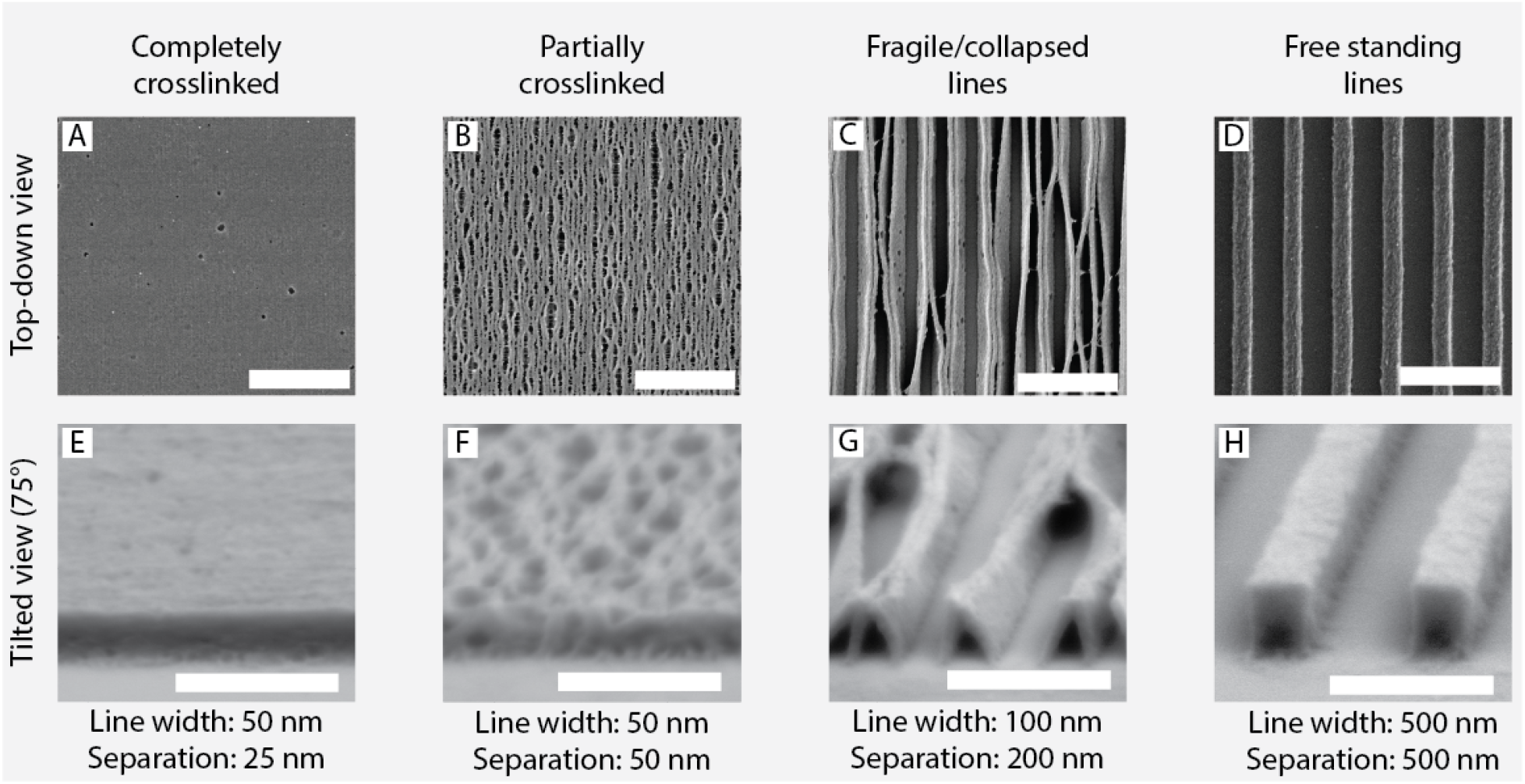
Line pattern fabrication outcome imaged using top-down and tilted view SEM. (A & E) Completely crosslinked pillar array where the individual lines are no longer possible to resolve. (B & F) Partially crosslinked lines where crosslinked SU-8 bring adjacent lines into contact. (C & G) Fragile/collapsed lines that contact each other or the substrate due to low stability. (D & H) Free standing lines, vertically aligned on the substrate. Scalebar 2000 nm for top-down view (A - D) and 1000 nm for tilted view images (E - H).

**Figure 7:**
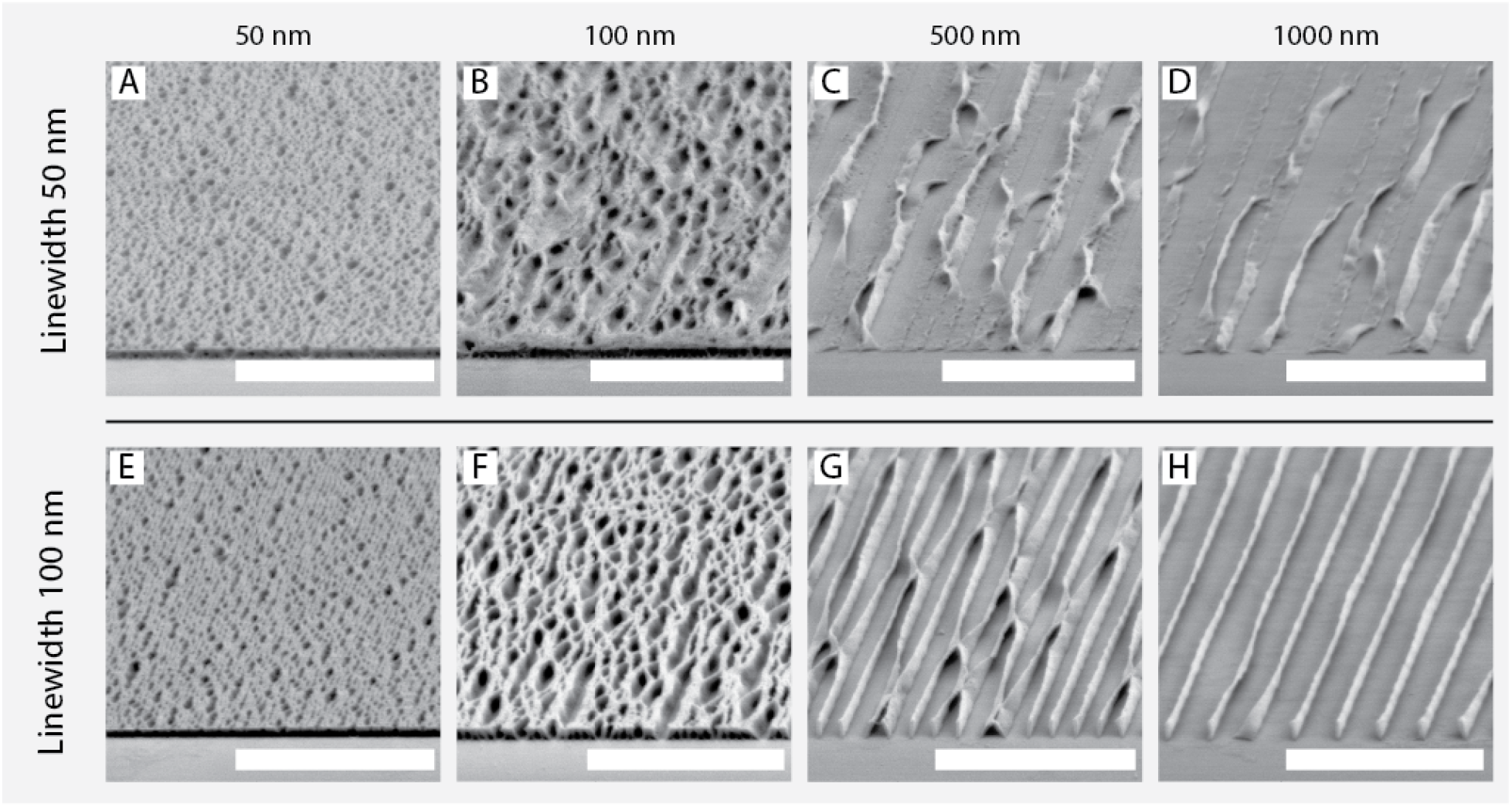
SU-8 lines with line width of 50 nm (A - D) and 100 nm (E - H) fabricated in a 500 nm resist with different separations between exposed lines. The thinner lines depicted in images (A - D) are fragile even at higher spacing, whereas the thicker (100 nm) lines depicted in images (E - H) become resolved and stable when the separation is increased. Indicating that the wider lines are able to overcome the capillary forces when sample is dried after development. Scalebar 5000 nm for all images.

When the line spacing was decreased, free single lines could no longer be produced. As indicated in Figure 5 too closely spaced lines resulted in either partial or complete cross-linking of the resist in the unexposed areas between the lines. The transition between free-standing/collapsed lines and cross-linking occurs at different separations depending on SU-8 film thickness. For thick resist films (2000 nm and 4000 nm) we were only able to fabricate lines with separations of approximately ten times the line widths, with a minimum line separation for 2000 nm films of 2000 nm and for 4000 nm separations of 2500 nm. For thin resists, it was possible to fabricate lines with separation as low as 50 nm (see Figure 5). In the intermediate thickness range, lines could be fabricated with widths of 50 nm (for 1000 nm SU-8 film) and 100 nm (for 500 nm SU-8 film and with separations of about 250 nm.

The fabrication outcome for 500 nm wide lines in different resist thicknesses is shown in Figure 8. When comparing similar lines exposed in SU-8 films at different film-thicknesses, we find that thinner SU-8 films allows for realisation of narrower and more closely spaced lines. For 500 nm wide lines, line separations of 250 nm was possible in 500 nm thick SU-8 films, but only 1000 nm and 2500 nm was possible in thicker films of 1000 nm and 2000 nm respectively.

**Figure 8:**
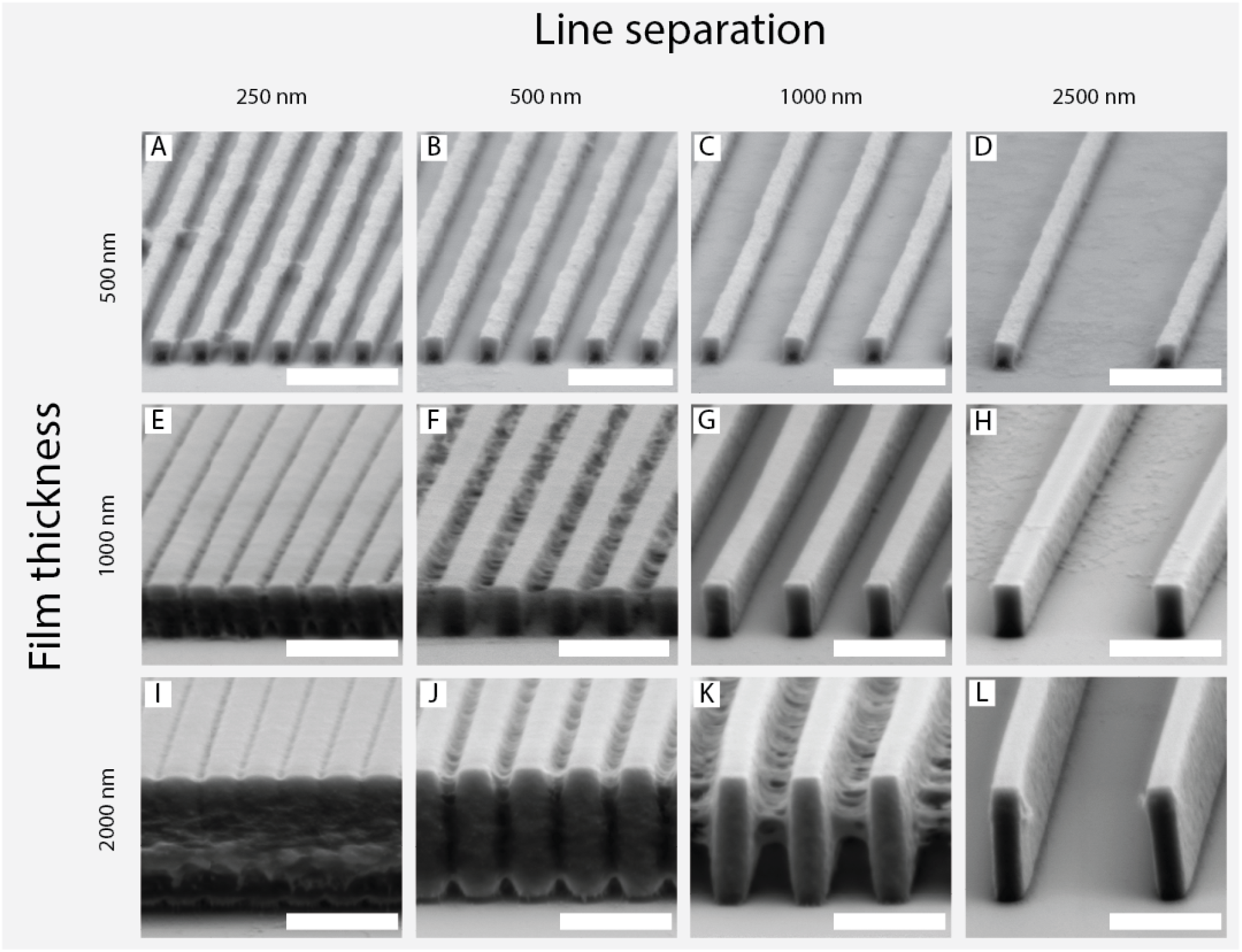
Fabricated line pattern, with 500 nm nominal line width, with variable line separation at three film thicknesses/structure heights. (A - D) Fabrication of lines in 500 nm thick SU-8 resist. Free standing lines are possible to fabricate down to 500 nm spacing. (E - F) Fabrication of lines in 1000 nm thick SU-8 resist, smallest possible spacing was 1000 nm. (I - L) Fabrication of lines in 2000 nm thick SU-8 resist, with a smallest possible spacing of 2500 nm. Due to structure broadening, the resulting line width after fabrication appears wider than designed, whereas the separation between the lines appear smaller. Micro-graphs taken at 75 degree tilt relative to the substrate. Scalebars 5000 nm for all micrographs.

The same optimisation was performed for nano-pillar arrays. The optimal dose for pillar arrays was found by testing a wide dose range for each combination of dot array pitch and SU-8 film thickness. Resulting surfaces were examined with SEM in top and tilted views. This exposure mode sets up a grid of dots and each dot is then exposed using the same dose. In contrast to the line pattern, no proximity effect correction (PEC) was performed for the dot arrays.

For grids of dots, no proximity effect correction is possible. In order to ensure a uniform dose throughout the pattern, we therefore exposed larger arrays of dots. When exposing 500 µm by 500 µm arrays of dots, the interior of the pillar arrays showed a uniform fabrication outcome. The imaging and classification of fabrication outcome was done in the centre area for all nanopillar arrays, except for Figure 11D, for where the pillar array edge was imaged for clarity.

In Figure 9 the fabrication outcome at the optimal dose for each array pitch is shown for all tested SU-8 film thicknesses. Electron dose that gave correctly reproduced pillars is noted for all combinations of film thickness and pillar pitches (Figure 9). In figure Figure 9 the fabrication outcome is colour coded. Green indicates fabrication of free standing pillars. In situations when the SU-8 cross-linking extended between the exposed dots, the outcome is colour coded with orange for partly crosslinked arrays and red for fully crosslinked arrays. The fabrication outcome for a given dot pattern is influenced by the electron dose and SU-8 film thickness. Below a critical dose, pillars with weak adhesion to the substrate are produced. The weak adhesion is attributed to the low surface area of the pillar at the base (see Figure 10A and E). At a threshold dose, pillars gain enough surface area to form free standing arrays. At this dose, fabricated structures have the highest possible aspect ratio, as any dose increase leads only to broadening and a lower aspect ratio (see Figure 10). The value of the critical dose depends on the resist thickness and the pitch of the fabricated array. Due to proximity effect, denser arrays have typically a lower threshold dose. As a result dot-arrays exposed with the same dose can produce different outcome for different pitches. Figure 10 shows the effect for 5000 nm, 1000 nm and 750 nm dot arrays in 2000 nm thick film. A 75 fC dose gives free-standing pillars for array with a pitch of 5000 nm, whereas it leads to partial cross-linking for arrays with the pitch of 1000 nm and 750 nm. For dot arrays with large separation between pillars, such as arrays with 5000 nm pitch, an increased dose leads to increased radial pillar dimensions.

**Figure 9:**
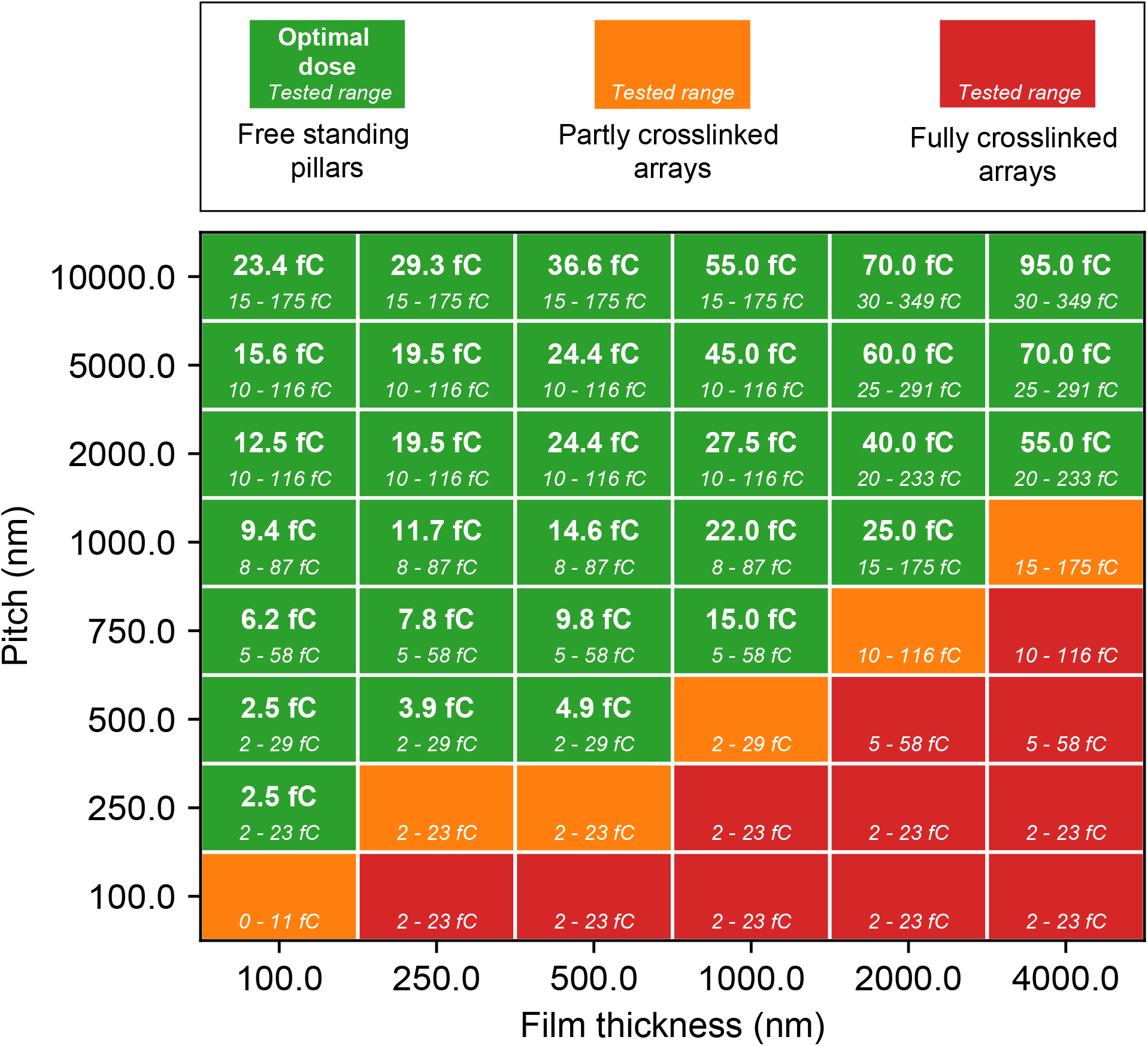
Colour coded representation of outcome of hexagonal pillar array fabrication with different film thicknesses and separation between exposed pillars. Free standing pillar arrays could be made with various thicknesses down to a pitch of 250 nm as described in Figure 3. In the cases where free standing pillars could be fabricated successfully, the working electron dose per dot/pillar is given. In italic font, the tested dose range for all combinations are noted.

**Figure 10:**
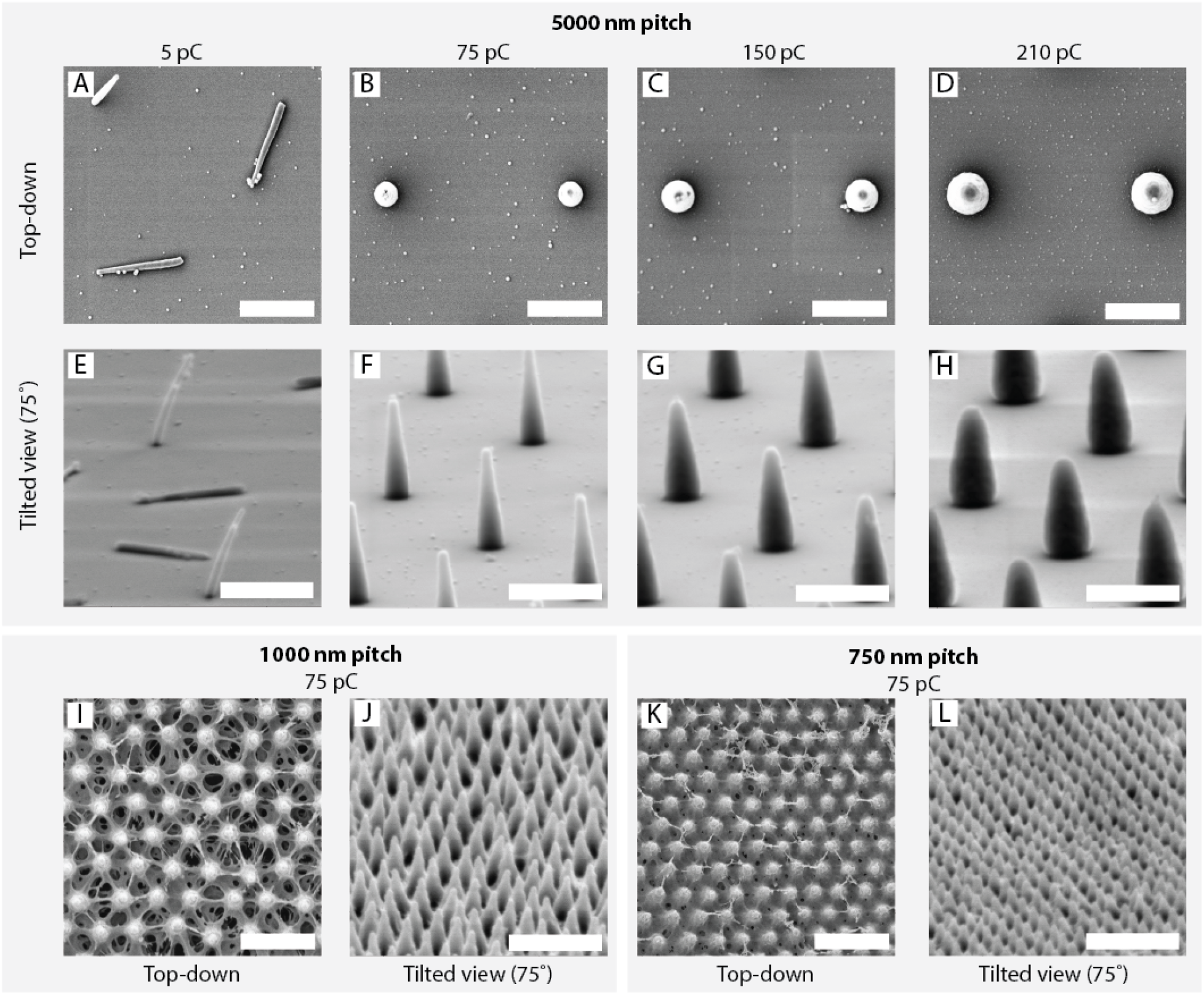
2000 nm high nano-pillars in SU-8 with 5000 nm pitch (images A - H), 1000 nm pitch (I and J) and 750 nm pitch (K and L) imaged in top-down and tilted view SEM. The pillars and the arrays are defined using the dot-pattern generator. (A & E) Below a given threshold the pillars are fragile and tend to fall over. (B & F) At an intermediate dose, pillars are stable and free standing. (C & G) For arrays with large separation between pillar locations, an increased electron dose leads to broader pillars. (D & H) Further increase in dose leads to even broader pillars. Using this effect, the pillar diameter can be controlled precisely. (I & J) 1000 nm pitched pillar array exposed with the same dose that lead to free standing pillars in the high-separation array. At this lower separation, the same dose gives partially crosslinked arrays. (K & L) When spaced even closer together, the trend continues, leading to more cross-linking at the same dose. Scalebars 2000 nm for all micro-graphs.

**Figure 11:**
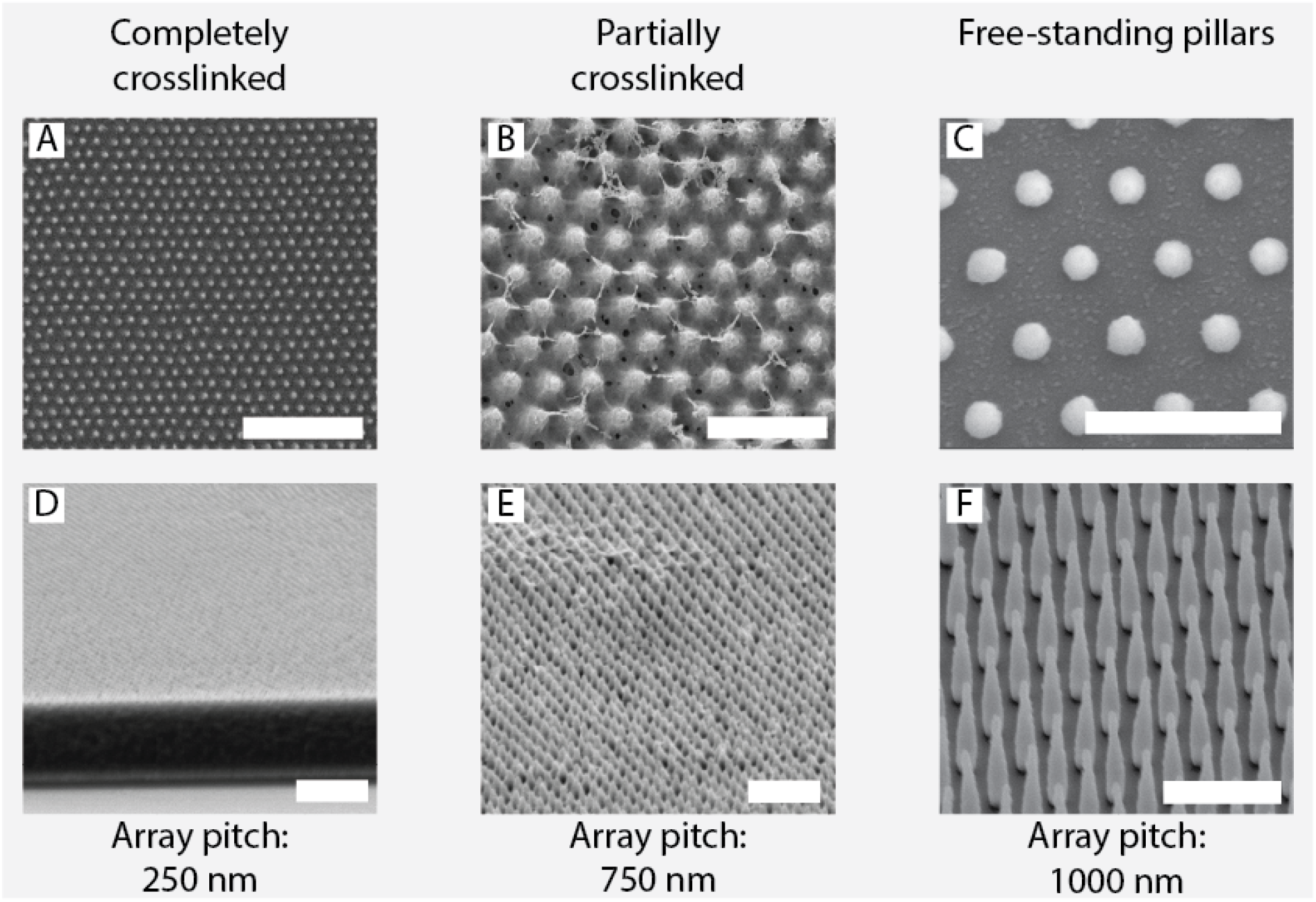
Scanning electron microscopy in top-down and tilted view was used to inspect the fabrication outcome for pillar fabrication using dot arrays. (A & D) When the pillar separation is very low, scattering causes the energy contrast between pillar location and areas in-between to be low and completely crosslinked arrays are formed. The position of the exposed dot is still visible in top-down view. (B & E) Partially cross-linked pillars where there is still SU-8 present between the locations of exposure, but the contrast in electron energy deposition is larger than for the completely crosslinked arrays. These arrays typically look rough, with the pillar tips constituting a rough surface. (C & F) If the array pitch, relative to the SU-8 film thickness was big enough, free standing arrays of vertically aligned SU-8 pillars could be formed. Scalebars correspond to 2000 nm for all micrographs.

When pillar arrays were fabricated without any adhesion promoter, pillar arrays exposed with a dose higher than the optimal dose showed irregular pillar detachment across the pillar array. In particular this was observed for low separation arrays at intermediate SU-8 film thicknesses. To increase the adhesion of SU-8 structures to substrates, a layer of HMDS was deposited by vapour deposition. Samples were placed in a desiccator together with vials of HMDS and left under HMDS atmosphere for 60 min. Pillar arrays exposed on samples with HMDS had a higher fraction of standing pillars compared to the samples without the adhesion layer. However, the application layer also influenced other aspects of processing, such as making it necessary to perform exposure and development straight after the resist spin coating.

For the thin SU-8 films (100 nm and 250 nm) it is possible to fabricate pillars with a minimum separation of about 2 times the resist thickness. In the intermediate SU-8 thickness (500 nm and 1000 nm) the minimum pitch is approximately the same as resist thickness, whereas in the thickest SU-8 tested (2000 nm and 4000 nm) the minimum separation between pillars is approximately half of the film thickness.

The exposure time for pillar arrays was minimised by using the dot-exposure writing mode. A 1 mm by 1 mm area of pillars with a spacing of 1000 nm could be exposed in 30 s, giving a writing speed of around 38000 pillars/s. If the pillars were exposed in the same way as lines were exposed in this work, the exposure of the same pattern with the same current would take an estimated 300 s mm*^−^*^2^.

To further characterise fabrication of the pillar arrays, we analysed the diameter of single pillars in the arrays at two locations, measured approximately 100 nm from the pillar top and about 100 nm from the pillar base. The “base diameter” is measured at the height of the pillar where the diameter is the biggest, as the pillars tend to taper slightly in towards the base. Results are summarised in Figure 12, where the mean value for tip and base diameter for all film thicknesses are plotted.

**Figure 12:**
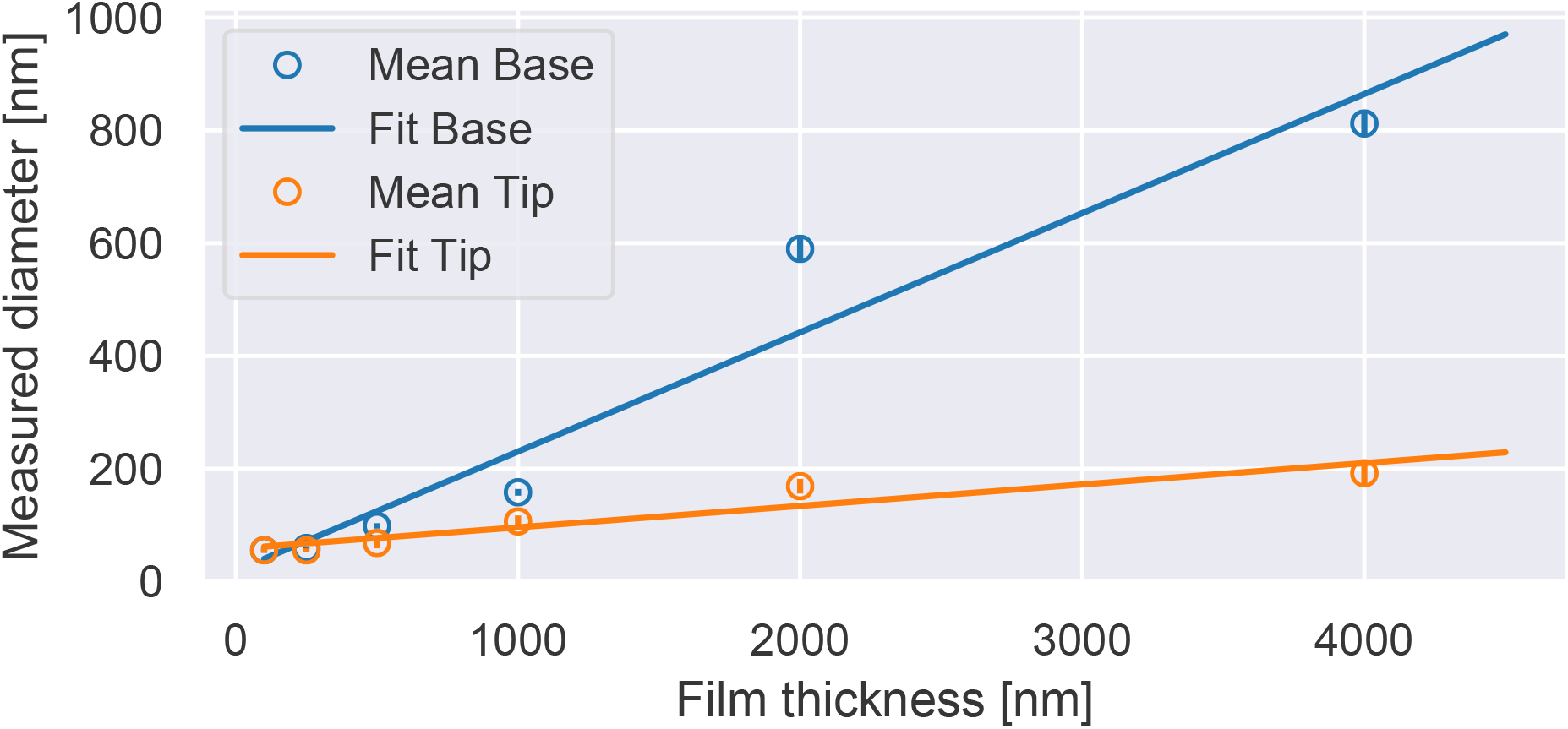
Measured pillar diameter close to the tip and close to the base of SU-8 pillars fabricated in films with various thickness. Mean value from six measurements are reported together with a the standard deviation and a linear fit to the mean value.

### 2.3 Topographies with multiple heights

Our approach can also be used to fabricate topographies with two distinctive heights. We demonstrate this by making nano-pillar arrays with heights of 3000 nm and 1000 nm. This is achieved by a process, where a second layer of SU-8 is spin coated onto a substrate containing a SU-8 resist that has already been exposed in EBL and subjected to a first post exposure bake. Without developing, a second layer of SU-8 can be spin coated. The sample is then soft-baked to remove solvent from the second resist layer. After the second exposure and and the second post exposure bake, both resist layers are developed simultaneously. Figure 13 shows a typical fabrication outcome. Pillars with the height of 3000 nm and 1000 nm spaced by 10 000 nm and 2000 nm respectively can be seen. Repeated processing has not affected pillar geometry post exposure bake, no increased cross-linking between structures defined in the two exposures is observed.

**Figure 13:**
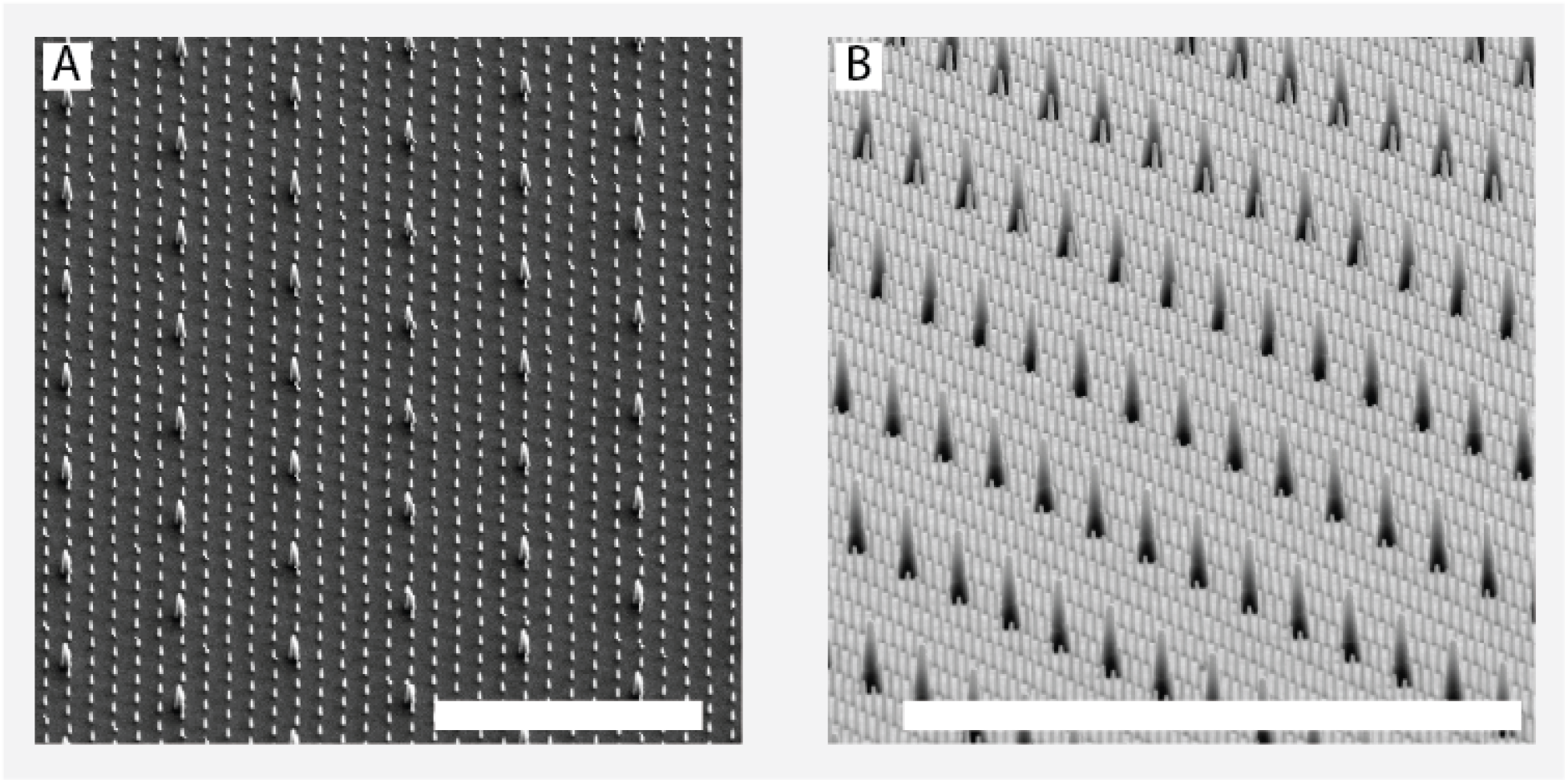
Two height hexagonal SU-8 pillar arrays superimposed onto each other. The lowest pillars are 1000 nm high with an array pitch of 2000 nm, whereas the higher pillars are 3000 nm with a separation of 10 000 nm. Scalebars 20 000 nm.

The second layer layer of SU-8 is spun on the first layer of structures without developing the first layer structures in between. By omitting the first layer development, the more viscous SU-8 can be spin coated easily. On the other hand, if the second layer of SU-8 is spin coated on developed SU-8 structures the structures cause a non-uniform second layer. In addition, the high aspect ratio SU-8 structures from the first layer can be detached from the surface.

Precise positioning of the two-layered structures require alignment of the layers relative to each other. By adjusting the position of the pattern relative to known reference features, over-layer alignment can be performed within a necessary tolerance (down to *≈* 10 nm for our EBL system). Alignment can be performed by using predefined reference, such as lithographically defined metal marks or as in the case of Figure 13 by using the corners of the squared sample.

To check whether the fabrication outcome was similar on another substrate, parts of the fabrication was also performed on Silicon in addition to the tests performed on glass substrates. Both of the test patterns were exposed and both possible fabrication as well as the optimal dose values working for glass also worked for SU-8 structures on Silicon, see Supporting Information.

### 2.4 Discussion

The present study was designed to establish the relation between resist thickness and the minimum size and separation of nano-structures that can be made from SU-8 resist on glass substrates using EBL. For the line patterns, we were able to fabricate 25 nm wide lines in 100 nm SU-8 films. The smallest possible width increased up to 250 nm for 4000 nm thick SU-8. For nano-pillar arrays, we found that the minimum possible separation between two pillars in an hexagonal array was 250 nm for 100 nm thin SU-8, increasing up to 2000 nm for 4000 nm.

Maximum structure height is limited by the electron scattering in the resist, and for 100 keV, our data is in agreement to what has been observed before.^30^ Structures with heights of 4000 nm to 5000 nm and a aspect ratio of 10 can be routinely fabricated. Broadening of the fabricated structures towards the base due to electron scattering in the resist in inherent for the EBL fabrication process. This leads to tapering of the structures, with the base diameter increasing greatly for thicker resist films, but with less increase in tip diameter.

Our results show that the high electron sensitivity of SU-8 resist facilitates high fabrication throughput without changing the exposure current. However, high sensitivity and low contrast makes the fabrication process more susceptible to proximity effects and limits the minimum separation between structures and the minimum feature size. Proximity effect correction (PEC) allows to reduce the proximity effect for line patterns by adjusting the dose throughout the pattern. For our test patterns, the effect from PEC is rather small. However for other pattern designs with larger differences in pattern density and feature size would probably see a larger improvement from the use of PEC.

For nano-pillar arrays that were written as single dots, PEC was not possible. Here, the fabrication process was instead altered in order to correctly reproduce pillar arrays in the central part of the large nano-pillar array. Proximity effects are influencing the fabrication outcome at the array boundary, where a region with a size in the range of 20 µm from the edge is underexposed resulting in collapsed or unstable pillars. In either of the fabrication approaches, the minimum feature size and minimum spacing is limited by the high sensitivity and low contrast of the SU-8.^31, 44^

Inspection of the fabricated pillars reveal no distorted pillars. Cross-linked pillars seem to have enough mechanical stability to withstand for example capillary forces arising when drying the samples after development. SU-8 is known to have good adhesion to Silicon substrates, but lower adhesion to glass.^33^ Preliminary experiments showed that indeed the adhesion to glass was a limiting factor for the fabrication, and HMDS was therefore added as an adhesion promoter between the glass and the SU-8. Including an adhesion promoter in the fabrication increased the adhesion, while it also influenced other aspects of the processing. If waiting steps were added between spin coating and development, arrays with low separation between pillars could no longer be realised as only completely crosslinked arrays could be produced. We speculate that this is a result of products (such as NH ^+^) from the silylation reaction of glass with HMDS slowly diffusing into the SU-8 and cross-linking it close to the substrate.^46^ We speculate that this contribution is already enough to partly cross-link the SU-8 due to high sensitivity and contrast of the resist.

Fabrication of SU-8 structures with multiple heights was possible by incorporating several coating/exposure steps and developing the layers simultaneously. In our exposures, we could fabricate structures in a second layer of SU-8 without influencing the already exposed structures. Structures exposed in both layers of SU-8 did not appear to be mechanically weaker than structures exposed in single-layer SU-8.

## 3 Conclusion

Nano-fabrication plays an important role in development of new tools for biomedical research. To support this development, 2D and 3D nano-fabrication approaches with the capability of fabricating high-resolution nanostructures with relatively high throughput is necessary. One method for realising such systems is electron beam lithography fabrication of vertically aligned surface bound nanostructures in the bio-compatible epoxy based polymer SU-8. This approach is capable of producing a range of different structure heights as well as in-plane resolution down to sub-100 nm directly on transparent substrates integrateble into common analysis techniques used in biomedical research. By bridging the gap between high resolution and high-throughput nano-fabrication, we envision that SU-8 nano- and microstructures fabricated with EBL will be a cornerstone in further development the field of biomedical research.

