## Supporting information for "Electron beam lithography fabrication of SU-8 polymer structures for cell studies"

#### Contents

**1 Comparison between exposure with and without PEC** **S2**

**2 Comparison between fabrication on glass and Si** **S2**

### 1 Comparison between exposure with and without PEC

To assess the influence from proximity effect correction on our line patterns, we exposed the same line pattern with and without proximity effect correction for given SU-8 film thicknesses. The outcome from these tests is presented and discussed here.

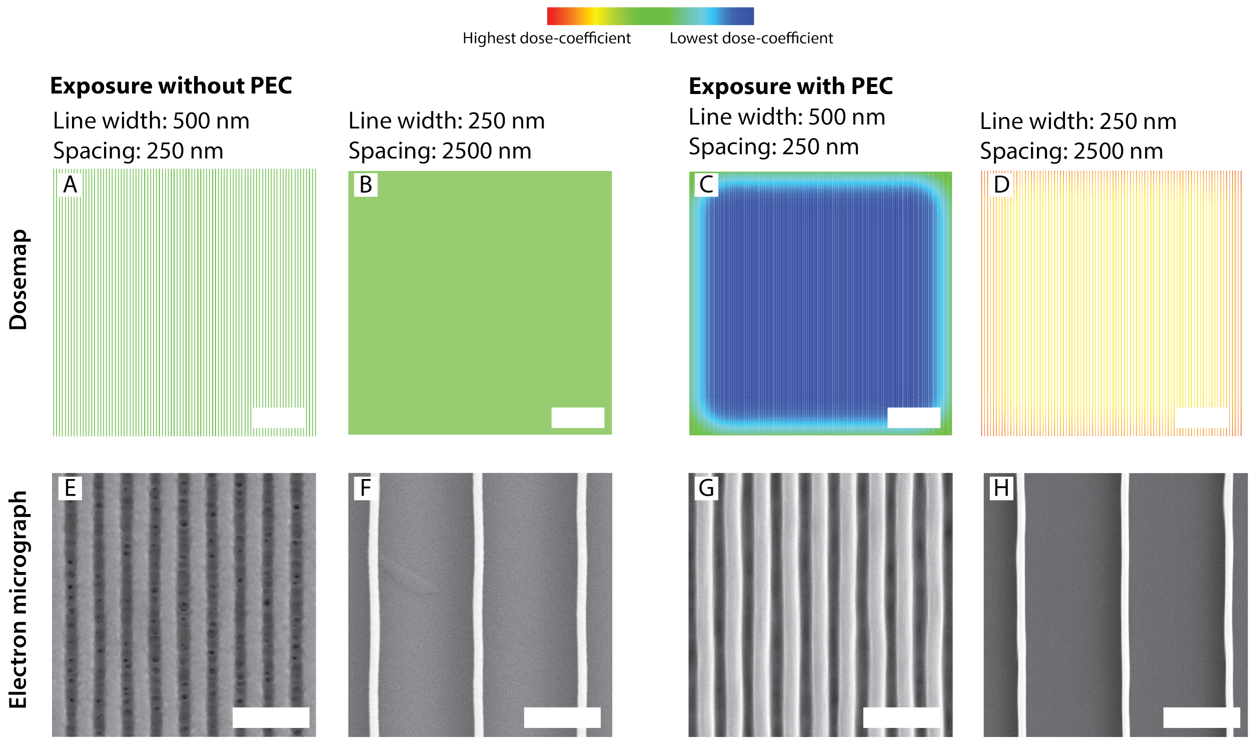


Figure S1: Dosemap showing dose-coefficients for 500 nm lines with 250 nm and 250 nm lines with 2500 nm spacing. For each of the cases, electron micrographs of the optimal fabrication outcome exposed with the same base dose for both the wide and narrow lines are shown. For the exposure without PEC, it is seen that the closely spaced lines are overexposed when exposed with the dose that is optimal for the narrow lines with high spacing. In the case of an proximity corrected exposure, the dose is adjusted in such case, that in most cases the same base dose can be used for both lines with high and low separation. The variations in pattern density and size is accounted for by the PEC. Scalebars: (A-D) 50 µm, (E-H) 2 µm.

### 2 Comparison between fabrication on glass and Si

In order to compare the processing conditions on glass and Silicon, we compared the processing conditions on glass and Silicon. The processing on Silicon was slightly altered, omitting the HMDS treatment as not adhesion problem between Silicon and SU-8 was needed in initial experiments.

**Fabrication SU-8 structures on glass** 24 mm x 24 mm glass cover slips (#1.5, Menzel-Gläser, 170 µm) were cleaned by immersion in acetone, isopropanol, and rinsed in de-ionised water followed by 2 min oxygen plasma treatment (Diener Femto plasma cleaner, power 100 W and pressure 0.4 mbar). The substrates were then dehydrated at 150 °C on a hotplate for 10 minutes. Hexamethyldisilazane (HMDS) was used as a adhesion promoter. It was applied by placing dehydrated glass substrates in a desiccator connected to a diaphragm pump and containing a vial of HMDS (Product Number: 440191, Sigma Aldrich). The sample was then kept under HMDS atmosphere for 60 min. Spin coating of the resist was performed straight after HMDS treatment.

SU-8 2001 was spin coated at 6000 rpm, soft-baked for 60 s and exposed in the Elionix EBL with a 500 pA beam current. Dose tests for line exposures were performed with base doses from 2 $\mu C\,cm^{-2}$ to 14 $\mu C\,cm^{-2}$ corrected using proximity effect correction as described previously. Pillar exposures were performed with the same dose ranges as described in the manuscript.

**Fabrication SU-8 structures on Si** 24 mm x 24 mm Si wafer (250 µm thickness) pieces cut from 2 inch Si (100) wafers were cleaned by immersion in acetone, isopropanol, and rinsed in de-ionised water followed by 2 min oxygen plasma treatment (Diener Femto plasma cleaner, power 100 W and pressure 0.4 mbar). The substrates were then dehydrated at 150 °C on a hotplate for 10 minutes. Spin coating of the resist was performed straight after dehydration bake.

SU-8 2001 was spin coated at 6000 rpm, soft-baked for 60 s and exposed in the Elionix EBL with a 500 pA beam current. Dose tests for line exposures were performed with base doses from 2 $\mu C\,cm^{-2}$ to 14 $\mu C\,cm^{-2}$ corrected using proximity effect correction as described previously. Pillar exposures were performed with the same dose ranges as

Table S1: Comparison between found optimal doses for pillar exposure in 1 µm thick SU-8 on Si and glass substrates.

| Pitch | Substrate material | |
|---|---|---|
| | Glass | Si |
| 100 nm | - | - |
| 250 nm | - | - |
| 500 nm | - | - |
| 750 nm | 15 fC | 10 fC |
| 1000 nm | 22 fC | 15 fC |
| 2000 nm | 27.5 fC | 20 fC |
| 5000 nm | 45 fC | 25 fC |
| 10 000 nm | 55 fC | 30 fC |

described in the manuscript.

**Results and Discussion** The same SEM analysis as presented in the main text was performed also to analyse the outcome from fabrication on Silicon substrates. In Table S1 and Table S2 the found doses are presented and compared to the optimal doses found on glass substrates.

For line patterns, the found base dose is then corrected by a pre-factor for each location in the pattern depending on the PEC.

Table S2: Comparison between found optimal base-doses for line-test pattern exposure in 1 μm thick SU-8 on Si and glass. The base doses were then corrected by the dose-coefficient from the proximity effect correction calculations.

| Line width | Separation between lines | Substrate material | |
|---|---|---|---|
| | | Glass | Si |
| 1000 nm | 10x (10 000 nm) | 7.0 μC cm$^{-2}$ | 5.0 μC cm$^{-2}$ |
| | 5x (5000 nm) | 7.0 μC cm$^{-2}$ | 5.0 μC cm$^{-2}$ |
| | 2x (2000 nm) | 10.5 μC cm$^{-2}$ | 5.0 μC cm$^{-2}$ |
| | 1x (1000 nm) | 10.5 μC cm$^{-2}$ | 5.0 μC cm$^{-2}$ |
| | 0.5x (500 nm) | 10.5 μC cm$^{-2}$ | 5.0 μC cm$^{-2}$ |
| 500 nm | 10x (10 000 nm) | 10.5 μC cm$^{-2}$ | 5.0 μC cm$^{-2}$ |
| | 5x (5000 nm) | 10.5 μC cm$^{-2}$ | 5.0 μC cm$^{-2}$ |
| | 2x (2000 nm) | 10.5 μC cm$^{-2}$ | 5.0 μC cm$^{-2}$ |
| | 1x (1000 nm) | 10.5 μC cm$^{-2}$ | 5.0 μC cm$^{-2}$ |
| | 0.5x (500 nm) | 10.5 μC cm$^{-2}$ | 5.0 μC cm$^{-2}$ |
| 250 nm | 10x (2500 nm) | 10.5 μC cm$^{-2}$ | 5.0 μC cm$^{-2}$ |
| | 5x (1250 nm) | 15.8 μC cm$^{-2}$ | 5.0 μC cm$^{-2}$ |
| | 2x (500 nm) | 15.8 μC cm$^{-2}$ | 8 μC cm$^{-2}$ |
| | 1x (250 nm) | 15.8 μC cm$^{-2}$ | 8 μC cm$^{-2}$ |
| 100 nm | 20x (2000 nm) | 10.5 μC cm$^{-2}$ | 5.0 μC cm$^{-2}$ |
| | 10x (1000 nm) | 10.5 μC cm$^{-2}$ | 5.0 μC cm$^{-2}$ |
| | 5x (500 nm) | 10.5 μC cm$^{-2}$ | - |
| 50 nm | 20x (1000 nm) | 10.5 μC cm$^{-2}$ | 5.0 μC cm$^{-2}$ |
| | 10x (500 nm) | 10.5 μC cm$^{-2}$ | - |
| | 5x (250 nm) | 10.5 μC cm$^{-2}$ | - |
